# Whole-genome sequence analysis unveils different origins of European and Asiatic mouflon and domestication-related genes in sheep

**DOI:** 10.1101/2021.09.07.458675

**Authors:** Ze-Hui Chen, Ya-Xi Xu, Xing-Long Xie, Dong-Feng Wang, Diana Aguilar-Gómez, Guang-Jian Liu, Xin Li, Ali Esmailizadeh, Vahideh Rezaei, Juha Kantanen, Innokentyi Ammosov, Maryam Nosrati, Kathiravan Periasamy, David W. Coltman, Johannes A. Lenstra, Rasmus Nielsen, Meng-Hua Li

**Affiliations:** CAS Key Laboratory of Animal Ecology and Conservation Biology, Institution of Zoology, Chinese Academy of Sciences (CAS), Beijing, China; University of Chinese Academy of Sciences (UCAS), Beijing, China; College of Animal Science and Technology, China Agricultural University, Beijing, China; Center for Computational Biology, University of California at Berkeley, Berkeley, CA 94720, USA; Novogene Co., Ltd, Tianjin, China; Department of Animal Science, Faculty of Agriculture, Shahid Bahonar University of Kerman, Kerman, Iran; Natural Resources, Natural Resources Institute Finland (Luke), Jokioinen, Finland; Board of Agricultural Office of Eveno-Bytantaj Region, Batagay-Alyta, Russia; Department of Agriculture, Payame Noor University, Tehran, Iran; Animal Production and Health Laboratory, Joint FAO/IAEA Division of Nuclear Techniques in Food and Agriculture, International Atomic Energy Agency, Vienna, Austria; Department of Biological Sciences, University of Alberta, Edmonton, AB T6G2E9, Canada; Faculty of Veterinary Medicine, Utrecht University, Utrecht, the Netherlands; Department of Integrative Biology, University of California at Berkeley, Berkeley, CA 94720, USA; Department of Statistics, UC Berkeley, Berkeley, CA 94707, USA; Globe Institute, University of Copenhagen, 1350 København K, Denmark

**Keywords:** *Ovis* genus, introgression, domestication, adaptation

## Abstract

The domestication and subsequent development of sheep are crucial events in the history of human civilization and the agricultural revolution. However, the impact of interspecific introgression on the genomic regions under domestication and subsequent selection remains unclear. Here, we analyze the whole genomes of domestic sheep and all their wild relative species. We found introgression from wild sheep such as the snow sheep and its American relatives (bighorn and thinhorn sheep) into urial, Asiatic and European mouflons. We observed independent events of adaptive introgression from wild sheep into the Asiatic and European mouflons, as well as shared introgressed regions from both snow sheep and argali into Asiatic mouflon before or during the domestication process. We revealed European mouflons arose through hybridization events between a now extinct sheep in Europe and feral domesticated sheep around 6,000 – 5,000 years BP. We also unveiled later introgressions from wild sheep to their sympatric domestic sheep after domestication. Several of the introgression events contain loci with candidate domestication genes (e.g., *PAPPA2*, *NR6A1*, *SH3GL3*, *RFX3* and *CAMK4*), associated with morphological, immune, reproduction or production traits (wool/meat/milk). We also detected introgression events that introduced genes related to nervous response (*NEURL1*), neurogenesis (*PRUNE2*), hearing ability (*USH2A*) and placental viability (*PAG11* and *PAG3*) to domestic sheep and their ancestral wild species from other wild species.

## Introduction

The genus *Ovis* spans ∼8.31 million years of evolution and comprises eight extant species: domestic sheep *O*. *aries*, argali *O. ammon*, Asiatic mounflon *O. orientalis*, European mouflon *O. musimon*, urial *O. vignei*, bighorn sheep *O. canadensis*, thinhorn sheep *O. dalli* and snow sheep *O. nivicola*^1^. Earlier archeological and genetic studies have provided strong evidence for that sheep have been domesticated from their wild ancestor Asiatic mouflon (*O*. *orientalis*) in the Fertile Crescent ∼12,000 – 10,000 years BP ^2–4^. The domestication during the Neolithic agricultural revolution had contributed significantly to human civilization by providing a stable source of meat, wool, leather and dairy.

In spite of varying diploid number of chromosomes (2*n* = 52 - 58) ^1^, hybridization between wild and domestic sheep, as well as between wild sheep species, has been documented to produce viable and fertile interspecific hybrids ^5–9^. Previous studies have shown genetic evidence for introgression ^10–14^, including adaptive introgression from wild relatives to domestic sheep ^15,16^. However, the importance of introgression in the entire *Ovis* genus and its contribution to the sheep domestication process remains largely unexplored.

Because wild sheep have adapted to different biogeographic ranges resulting in them being resilient to many biotic and abiotic stresses, the existing genetic variation of wild sheep provide an important genetic resource for improving domestic sheep in response to increased food production demands, animal disease occurrence and rapid global climate change. Elucidating the evolutionary and genetic connection between wild and domesticated sheep is therefore important for understanding the potential for using wild sheep genetic material for improvement of domesticated sheep.

In this study, we use high-depth whole genome sequences (average coverage = ∼21×) of 72 individuals from the eight *Ovis* species, most of which were understudied in previous genomic studies ^7,17–19^. We reconstructed the phylogeny and evolutionary history of these species. In addition, we explored gene flow between species and selection signatures of domestication. These findings add to our understanding of the origins of the Asian and European mouflons and the emergence of domestic sheep.

## Results

### Sequencing and variant calling

High-depth resequencing of 72 individuals from eight *Ovis* species (Fig. 1a and Supplementary Table 1) generated a total of 35.91 billion 150-bp paired-end reads (5.39 Tb), and 35.84 billion clean reads (5.28 Tb) with an average depth of 20.7×(12.2 – 36.9×) per individual and average genome coverage of 97.2% (96.5% – 98.3%) after filtering. The average sequence coverage was 19.3× for *O. aries*, 17.8× for *O. ammon*, 18.9× for *O. canadensis*, 19.8× for *O. vignei*, 18.9× for *O. musimon*, 17.8× for *O. nivicola*, 19.4× for *O. dalli*, and 27.1× for *O. orientalis.* On average, 95.83% individuals had ≥ 4× coverage, 90.11% had ≥ 10× coverage, and 46.68% had ≥ 20× coverage. Of all the individual sequencing reads, 91.86% were mapped to the *O. aries* reference genome Oar_v4.0 (Supplementary Table 2). Summed over all samples, 125,982,209 SNPs, 13,043,920 INDELs (insertions and deletions ≤ 50bp; ∼0.89 million common indels shared by all sheep species and on average 2,605,718 per individual) (Table 1 and Supplementary Tables 2 – 6) and genome-wide structural variations (SVs, 51bp – 997.369 kb: inversions, insertions, deletions, duplications and translocations, on average 41,965 per individual), including copy number variations (CNVs, deletions and duplications of 51bp to 997.369 kb, on average 31,124 per individual) (Supplementary Table 7) were detected. The number of SVs shared by two species ranged from 29,884 to 91,186 (Supplementary Table 8 and Supplementary Fig. 1a). On average, 1.88% SVs were located in exonic regions, 65.2% SVs were located in intergenic regions, and 29.9% SVs were located in intronic regions, while 67.0%, 31.0% and 0.66% SNPs were in intergenic, intronic and exonic regions, respectively (Supplementary Tables 9 and 10).

**Figure 1.**
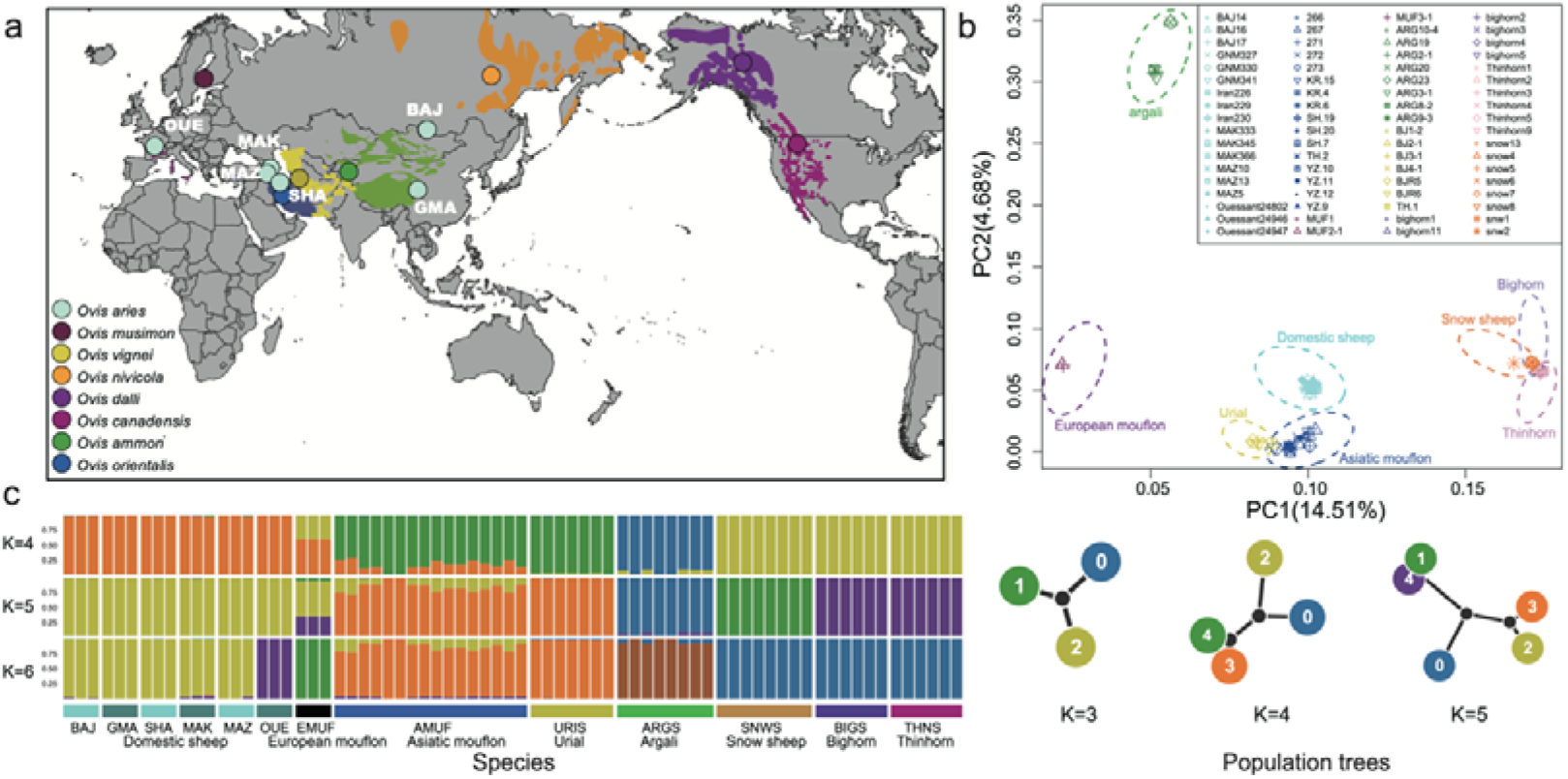
Geographic distribution and population structure of *Ovis* species. **(a)** Geographic map of sample location and wild sheep species distribution based on the IUCN Red list (https://www.iucnredlist.org). Here we adopted the classification of Nadler *et al*. 1973. **(b)** Principal Component Analysis (PCA) of *Ovis* species. **(c)** Admixture plot using Ohana software for K from 4 to 6. Population tree of each K indicates affinity of each ancestral component (Below, right).

**Table 1.** Summary information of whole-genome variations identified in *Ovis* species.

| Sequence quality/<br>Variation type | *O.aries*<br>(n=18) | *O.orientalis*<br>( n=17 ) | *O.musimon*<br>(n=3) | *O.vignei*<br>(n=7) | *O.ammon*<br>(n=8) | *O.nivicola*<br>(n=8) | *O.dalli*<br>(n=6) | *O.canadensis*<br>(n=6) |
| --- | --- | --- | --- | --- | --- | --- | --- | --- |
| SNPs | 31,535,487 | 53,618,832 | 13,360,034 | 30,125,980 | 25,160,871 | 22,845,295 | 23,185,374 | 23,069,044 |
| INDELs | 4,361,226 | 7,173,026 | 3,180,207 | 4,748,458 | 4,541,250 | 4,204,319 | 4,301,920 | 4,255,087 |
| SVs | 123,594 | 161,892 | 55,950 | 81,003 | 84,587 | 75,375 | 77,304 | 75,480 |
| CNVs | 74,672 | 92,366 | 37,250 | 56,216 | 52,236 | 48,705 | 48,794 | 48,527 |
| Duplications | 1,814 | 2,403 | 571 | 915 | 1,185 | 1,071 | 1,122 | 1,070 |
| Deletions | 72,858 | 89,963 | 36,679 | 55,301 | 51,051 | 47,634 | 47,672 | 47,457 |
| Insertions | 11,685 | 12,981 | 6,517 | 8,126 | 8,867 | 7,339 | 8,954 | 8,727 |
| Inversions | 746 | 1,011 | 311 | 499 | 600 | 583 | 580 | 576 |
| Translocations | 36,491 | 55,534 | 11,872 | 16,162 | 22,884 | 18,748 | 18,976 | 17,650 |
| Average Depth (X) | 19.25 | 27.11 | 18.89 | 19.77 | 17.78 | 17.8 | 19.43 | 18.88 |
| Coverage Rate (%) | 97.2 | 98.22 | 97.03 | 97.1 | 96.79 | 96.54 | 96.65 | 96.62 |

The percentage of SNPs that was present in public databases [e.g., NCBI sheep dbSNP database v150 and European Variation Archive (EVA)] ranges from 78.1% in argali to 94.3% in domestic sheep (Supplementary Table 11). Our dataset added 2,139,962 novel SNPs (an increase of 7.04%) to the NCBI and EVA database of sheep genetic variants (Supplementary Table 6). Of the 176,403 common sites between detected SNPs and the Ovine BeadChip, an average of 288,638 genotypes observed here were validated by the Ovine Infinium HD SNP BeadChip data available for 14 individuals of the samples sequenced (97.1% validation rate, and 297,115 common SNPs), and an average of 23,220 genotypes were validated by the Ovine SNP50K BeadChip data available for another 12 individuals of the samples (96.64% validation rate, and 22,583 common SNPs) (Supplementary Table 11).

Moreover, 74 randomly selected SNPs, which are from the NCBI sheep dbSNP database and the candidate genes identified below, were inspected in 4-12 individuals by Sanger sequencing and produced an overall validation rate of 95.5% (Supplementary Table 12). For PCR and qPCR validation of CNVs (deletions and duplications), 14 randomly selected CNVs with 85.4% concordant genotypes (38/42 deletions and 32/40 duplications; Supplementary Table 13 and Supplementary Fig. 2) were successfully validated. The validation rates observed here are higher than those in previous studies^17,20^, which could be due to more efficient and precise CNV detection methods used here. The high validation rate indicated high reliability of the genetic variants created in this study.

### Patterns of variation

The 126 million SNPs were detected across all eight species. The number of SNPs varied from 11.3 to 20.1 million per individual and from13.4 to 53.6 million (0.6 – 18.2 unique) per species (Supplementary Table 2, Supplementary Table 6 and Supplementary Fig. 3c). We observed 4,431,063 SNPs shared among all the eight species, with the shared SNPs for pairwise comparisons varying from 6,241,176 between European mouflon and snow sheep to 25,195,033 between Asiatic mouflon and urial (Supplementary Table 4 and Supplementary Table 6). More comparisons of structural variants (SVs) and copy number variants (CNVs) among species for uniqueness and sharing are shown in Supplementary Fig. 1c.

Using pairwise genome-wide *F*_ST_, the species with highest genetic differentiation were snow sheep and European mouflon, and the ones with least differentiation were urial and Asiatic mouflon (Supplementary Table 14). The species with the highest genomic diversity (π), when only including SNPs with < 10% missing data, were domestic sheep, Asiatic mouflon, and urial (0.0032 – 0.0044), and the ones with lowest diversity were snow sheep, bighorn sheep and thinhorn sheep (0.00075 – 0.00078) (Supplementary Fig. 4b). On average, 67.0% of SNPs were located in intergenic regions, 31.0% in introns, and 0.7% SNPs in exons. The ratio of non-synonymous to synonymous substitutions ranged from 0.72 in urial and 0.77 in domestic sheep to 0.88 in European mouflon (Supplementary Table 10). We pooled the SVs and CNVs across all eight species yielding a high depth of coverage for the shared and unique SVs and CNVs among them (Supplementary Figs. 1a, b and Supplementary Tables 6 and 7). Annotation of genes overlapped with SVs were summarized in Supplementary Table 15 (see Supplementary Information)

### Phylogenomic reconstruction among the *Ovis* species

We generated eight high-depth whole pseudo-haploid genomes (see Online Methods), representing the eight *Ovis* species. Phylogenetic trees were then constructed from concatenated protein coding regions (CDSs) of autosomes, the X chromosome and the mitogenome of the assembled genomes, separately (Supplementary Fig. 5). These trees showed different phylogenetic patterns, but a consistent split between the three Pachyceriform species (i.e., bighorn, thinhorn and snow sheep) and the others, consistent with earlier genetic studies ^1,21^. Together with the observation that the first fossil evidence of caprinae is in the Upper Vallesian in Spain ^21^, these trees confirmed a Eurasian origin of the ovine species ^15,22^.

We split the whole genome (one high-depth genome per species, see Materials and Methods) into 2,462 autosomal and 136 X-chromosomal 1Mb non-overlapping windows of each species, and estimated Maximum likelihood (ML) trees for these windows. Three topologies (A, B and C) were observed for 46.1%, 29.1% and 17.8% of the autosomal trees, 33.8%, 50.0% and 7.4% of the X chromosomal trees, respectively (Supplementary Fig. 6). The main topologies A and B were also found using the maximum likelihood estimation on the concatenated CDSs (topology A, Fig. 2b) and using consensus methods of the Densitree on the non-overlapping fragments for autosomes (topology A, Fig. 2a) and X-chromosome (topologies B, Fig. 2a). We also estimated trees using high-depth individual autosomes and X-chromosome (Supplementary Fig. 5), which also support topologies A and B, respectively, while the individual mtDNA tree did not resemble any of the nuclear topologies.

**Figure 2.**
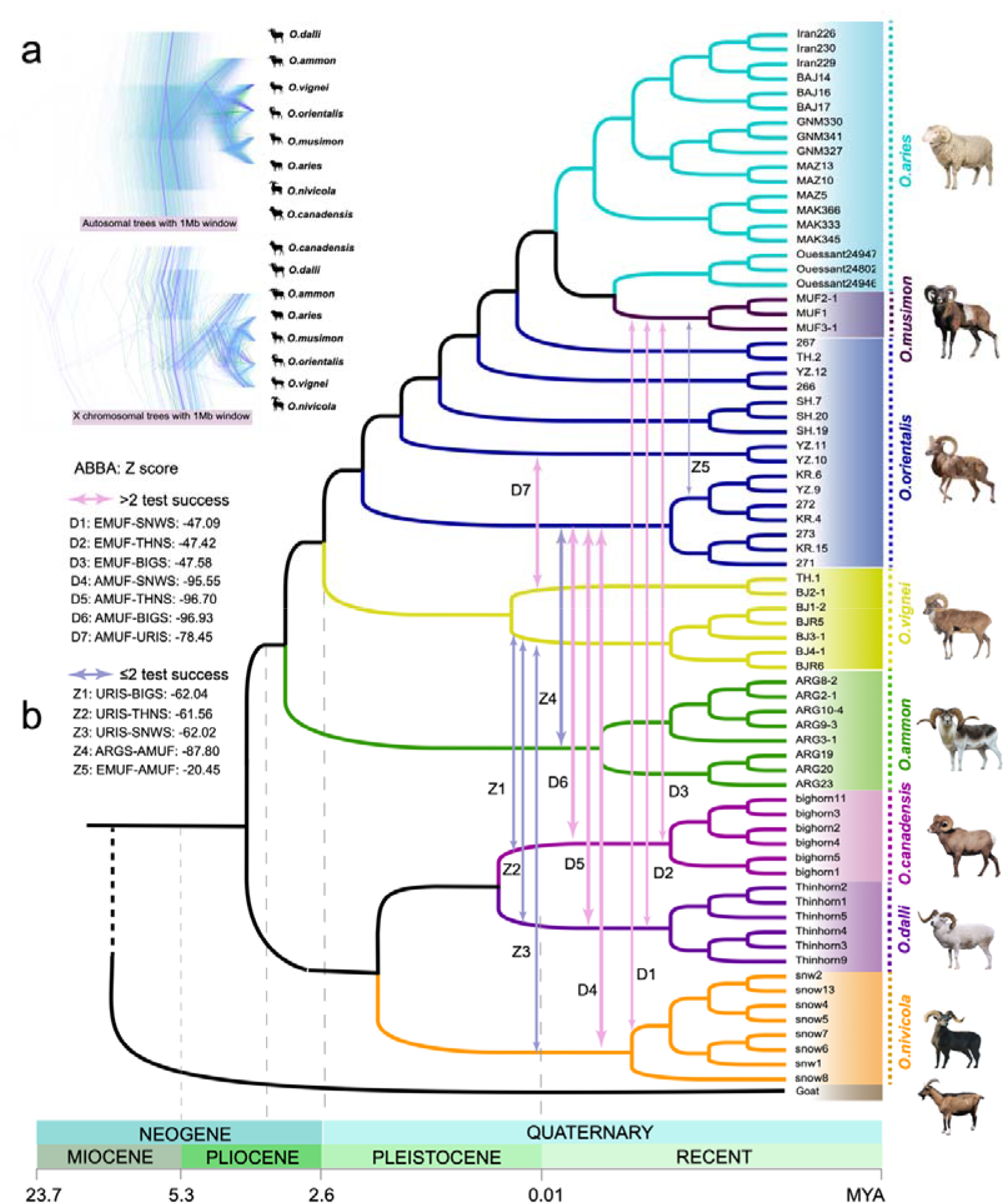
Phylogeny of *Ovis* genus. Prevalent discordance among segmental trees on autosomes and X chromosome, totaling 2,598 1-Mb segments. The segmental trees were visualized by Densitree. Consensus tree topologies of each category are shown in purple. **(b)** Phylogenetic tree of whole autosomal coding region (CDS) of 72 individuals using RAxML. Arrows marked the introgression pairs and the corresponding Z score based on the four populations test (*D* statistics), see Supplementary Table 17. Pink arrows indicate the admixed pairs which have been selected by three or more of the tests such as *D* statistics, TreeMix, *f* statistics and admixture analysis, while purple arrows indicate the introgression pairs detected in two or less of the tests.

The minor topologies (e.g., B and C for autosomes; and A and C for X-chromosome) may reflect local introgression between the species or incomplete lineage sorting (ILS) of ancestral phylogenies. The three phylogenies of all the 72 individuals using concatenated CDSs of autosomes (Fig. 2b and Supplementary Fig. 7a), X-chromosome and mitogenomes (Supplementary Fig. 7) showed seven major clades of individuals, with European mouflon sequences located among domestic sheep, compatible with the assumption that European mouflon is a domestic sheep subspecies ^15^. Also, European mouflon and domestic sheep show the same diploid number of chromosomes (2*n* = 54) ^1^.

The phylogenetic trees (Supplementary Figs. 5, 7) and pairwise *F*_ST_ (Supplementary Fig. 8b) showed two clear clusters, one comprised of the European mouflon and domestic sheep, and another of the Asiatic mouflon and the urial sheep. This is again compatible with the hypothesis that the European mouflon is a feral derivative of domestic sheep, but it also suggested that the Asiatic mouflons, sampled in Iran, have diverged considerably from the mouflon ancestors of both the early domestic hair sheep and the domestic wool sheep of more recent origin ^22^.

A coalescent hidden Markov model (CoalHMM) based on autosomal sequences indicated a divergence time of domestic sheep and the three Pachyceriform species of 0.244 to 0.270 mya. The argali and the urial were estimated to have diverged from domestic sheep ∼0.124 – 0.150 mya and ∼0.077 – 0.092 mya, respectively (Supplementary Figs. 9 – 11). The divergence time of the Asiatic mouflon and the urial was estimated to have occurred 0.073 – 0.083 mya, which is earlier than the divergence between bighorn and thinhorn sheep (∼0.036 – 0.052 mya). An Isolation with Migration (IM) model ^23^, which incorporates the impact of migration among species, gave a similar estimation with the Isolation (I) model ^23^.

The relatively recent divergence of the European mouflon from domestic sheep 5,550 – 5,450 BP (Supplementary Figs. 9 – 11) is concordant with the paleontological evidence of teeth and bone for a divergence of the Corsican mouflon and domestic sheep dated at 6,000 – 5,000 BP ^24^. Moreover, the coalHMM, IM and I models, with a filtering thresholds of < 1,000 years and > 20,000 years for the split time ^14^, showed a split time of 12,800 - 8,800 BP between domestic sheep and the Asiatic mouflon. This estimate is congruent with the estimated domestication time of sheep from the Asiatic mouflon around 9,000 – 11,000 BP, based on archaeological data ^4,24^, and also is in agreement with the time range 12,000 BP-8,000 BP from the start of exploitation to the end of domestication ^25^.

### Demographic history

The pairwise sequentially Markovian coalescent (PSMC) model found a dramatic decline in population sizes of these species ∼80 – 250 thousand years ago (kya) with a bottleneck for urial and Asiatic mouflon during 30,000 – 10,000 BP (Fig. 3a), coinciding with the glacial periods. The subsequent increase in their population sizes can be ascribed to the prosperity of animal husbandry, agriculture and sedentarism ^26^. The SMC++ analysis showed a decline of all species 10,000 – 1,000 BP. In particular, we noted European mouflon has a more dramatic decline of *N*e than domestic sheep 6,000 – 5,000 BP, which probably corresponds to the feralization of the European mouflon (Fig. 3c). The split between domestic sheep and the Asiatic mouflon occurred during 15,000 – 9,000 BP. During this time period, the Asiatic mouflon showed an increased *Ne*, whereas domestic sheep experienced a severe bottleneck because of domestication.

**Figure 3.**
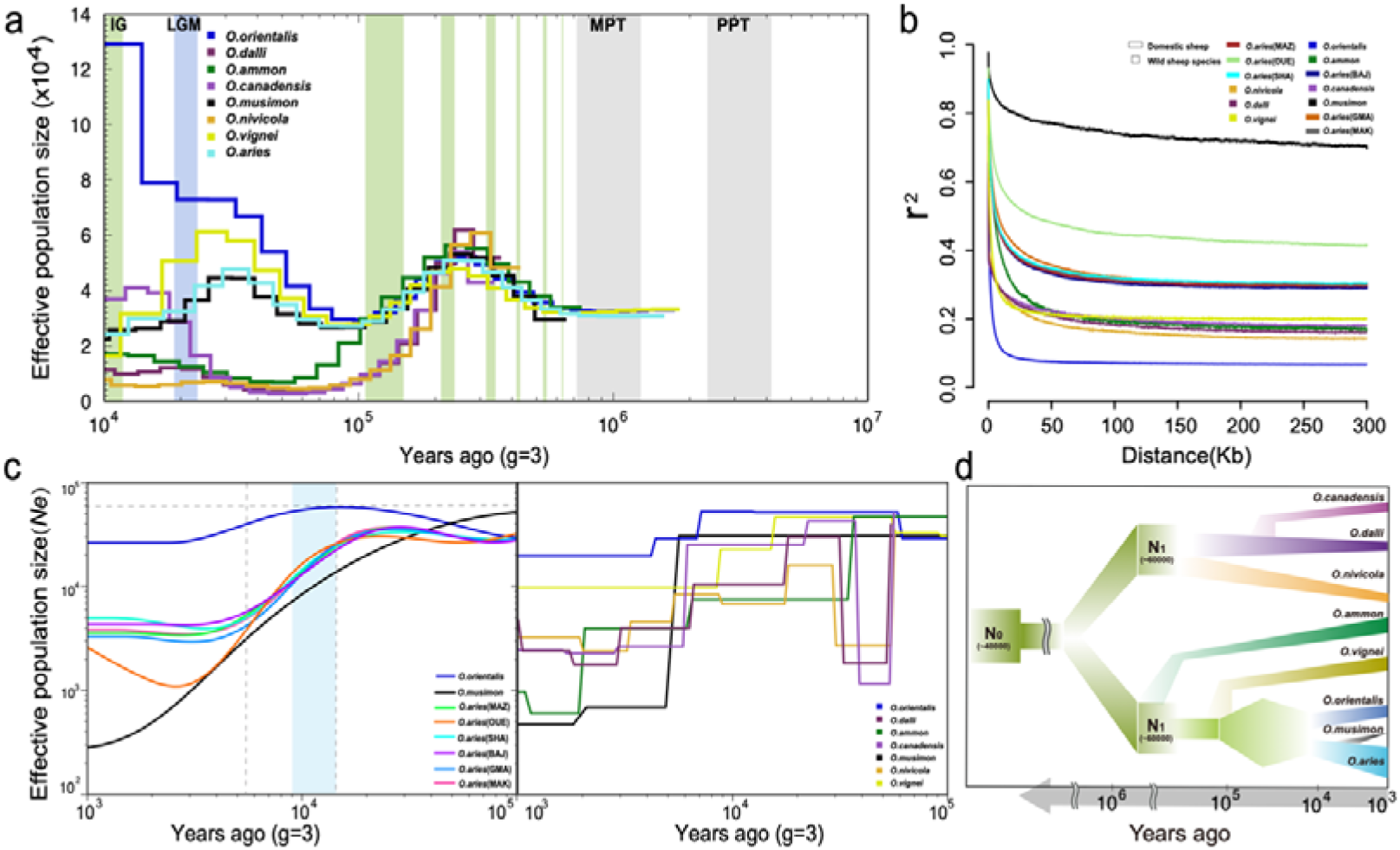
Demographic inference. **(a)** Ancestral dynamic change of effective population size inferred by PSMC program for eight high-depth genomes. Colors of the lines indicate different species. Plots were scaled using a mutation rate of 2.5×10^−8^ per site per generation and generation time (g) of 3. Light green shading indicates interglacials (IG) in the Pleistocene and Holocene, and light blue marked as LGM the Last Glacial Maximum and gray shading indicates the mid-Pleistocene transition (MPT) and the Plio-Pleistocene transition (PPT). **(b)** LD decay analysis for seven wild species (marked as rectangles) and six domestic breeds (marked as squares). **(c)** Dynamic change of effective population size inferred by the SMC++ program for Asiatic and European mouflon and six domestic breeds (left panel) and seven wild species (right panel), the blue shading indicates the period of domestication and the gray vertical dashed line is the potential split time point of European and Asiatic mouflon. **(d)** Dynamic change of effective population size over time for all *Ovis* species.

### Genetic structure and differentiation

PCA clusters individuals according to the recognized eight species. The cluster of argali showed significant within-species genetic divergence (Fig. 1b), which was also observed in the admixture pattern at high *K* values (Supplementary Fig. 12). The Asiatic mouflon cluster was dispersed and overlaps partially with the urial cluster (Fig. 1c and Supplementary Fig. 12). The population tree was compatible with the inferred genetic clustering at *K* =11 (Supplementary Figs. 12a, b), in which each species is assigned its own components. The admixture plot may suggest gene flow from argali (0.06 – 0.76%) and urial (1.4% – 15%) to Asiatic mouflon and possibly from wild relatives to domestic sheep, such as from European mouflon (5.4 – 5.7% of the genomes at *K*=6) to Ouessant sheep, which was an isolated island domestic breed (Supplementary Fig. 12). However, we noted that admixture proportions cannot be interpreted a direct evidence of admixture.

We observed higher levels of linkage disequilibrium (LD) in European mouflon and domestic sheep than in other species (Fig. 3b). This may be explained by a strong bottleneck during domestication. The Ouessant sheep ^27^ clearly had a higher LD than other domestic breeds, which was consistent with their low genomic diversity (Fig. 3b and Supplementary Fig. 4). Likewise, the high LD in European mouflon could be explained by a small population size and possible bottleneck during its reintroduction from Corsica island to continental Europe ^15^.

### Genomic introgression between wild species

The ABBA-BABA analysis (*D*-statistic) was implemented using ANGSD-based on alignments, which suggested introgressions from bighorn, thinhorn and snow sheep into their Eurasian relatives such as urial and Asian and European mouflon. (Supplementary Table 16). Statistical analyses based on variants using Admixtools (Supplementary Tables 17, 18), TreeMix (Supplementary Figs. 13 and 14) and *f*_d_ statistics (Supplementary Fig. 15 and Supplementary Table 19) consistently showed significant introgression of snow, bighorn and thinhorn sheep into urial, Asiatic and European mouflon. Bighorn and thinhorn sheep showed similar patterns of introgression as snow sheep in terms of several statistic indices, such as percentage (urial: 6.23 – 6.33%, Asiatic mouflon: 3.63 – 3.7%, European mouflon: 1.43 – 1.47%), length (urial: 152.68 –155.1Mb, Asiatic mouflon: 88.96 – 90.7 Mb, European mouflon: 35.08 – 35.96Mb) and shared genes (urial: 720 –744, Asiatic mouflon: 449 – 468, European mouflon: 151– 155) of introgression (Supplementary Fig. 15). For simplicity, we will focus only on the snow sheep introgression. The introgression events into urial and Asiatic mouflon had a lot of overlap in terms of genomic regions, while there was very minimal overlap between Asiatic and European mouflon introgression segments (Supplementary Fig. 15 and Supplementary Table 17). Furthermore, admixture graph fitting based on *f*_4_ statistics was carried out using the R package *admixturegraph* (Fig. 4), indicating a very close relationship between wild sheep of Pachyceriforms and European mouflon.

**Figure 4.**
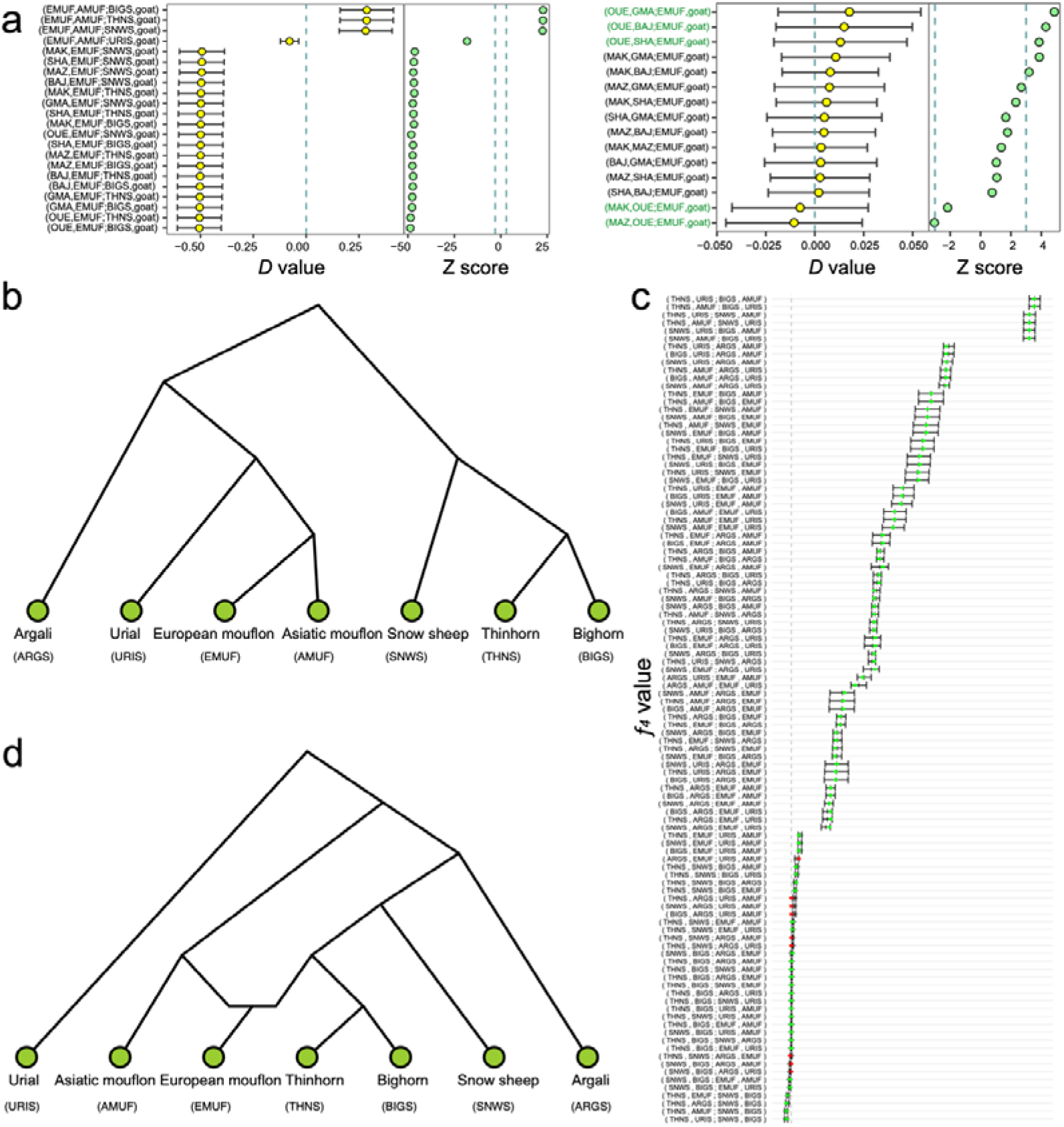
Admixturegraph fitting for introgression from the Pachyceriforms into European mouflon. **(a)** *D* statistics of European mouflon (EMUF) with the Pachyceriforms [snow sheep (SNWS), bighorn (BIGS) and thinhorn (THNS)]. **(b)** Prior phylogeny of wild species in *Ovis* genus. **(c)** Goodness of fit of *f*_4_ statistics. **(d)** Admixture graph.

Signatures of introgression were detected in candidate regions overlapping 892 genes from snow sheep to urial sheep, these genes were significantly (False Discovery Rate, FDR of 0.05 by the method of Benjamini-Hochberg^28^) enriched for nerve conduction, energy metabolism, membrane signal transduction, bile secretion, drug addiction and motor activity using DAVID annotation tools. From snow sheep to Asiatic mouflon or European mouflon, we found candidate introgression regions covering 497 and 179 genes, respectively (Supplementary Fig. 15a). In European mouflon, the introgressed genes were enriched for nerve regulation, locomotory behavior, cardiac disease, insulin secretion, serotonin metabolic process and calcium signaling pathway, while in Asiatic mouflon the genes were enriched in walking behavior, regulation of cell differentiation, ovarian steroidogenesis and platelet activation. Noteworthy, we observed three shared GO terms for the genes involved in the inter-species introgression events, such as motor, iron channel activity, and dendrite development. (Supplementary Table 20).

Among the three sets of introgressed genes between wild species, we observed 12 shared genes (*CYP2J, PRUNE2, ZNF385B, IMMP2L, GRIK2, HS6ST3, USH2A, LOC101111335, TMEM132D, PAG11, PAG3* and *CTNNA3*), which have functions associated with reproduction and production traits such as follicular development (*CYP2J, IMMP2L*), prolificacy (*GRIK2*), growth (*HS6ST3*), wool and body weight (*TMEM132D*) ^29–34^, and nervous response such as hearing ability evolution (*USH2A*) and nerve development (*PRUNE2*) ^35,36^. In particular, shared signatures of introgression were observed in the PAG gene family, which is involved in pregnancy detection and placental viability evaluation ^37^. Moreover, these genes were significantly enriched in a GO term (GO:0004190), which consists of pregnancy-associated glycoproteins (*PAG3* and *PAG11*) related to aspartic-type endopeptidase activity. We also observed one marginally significantly enriched KEGG pathway of protein digestion and absorption (oas04974) including the two genes from the PAG gene family.

### Dating the introgression from wild relatives to Asiatic mouflon

In addition to the introgression from snow sheep to Asiatic mouflon (Figs. 5a – d) mentioned above, *D*-statistics, *f*_3_ statistics and TreeMix analysis also detected signatures of introgression from argali into Asiatic mouflon (Supplementary Tables 16 – 21 and Supplementary Figs. 13 and 15a). Across-genome *f*_d_ values detect 670 and 734 segments introgressed by snow sheep and argali, respectively, corresponding to a genomic coverage of 3.68% and 3.98%, containing 497 and 540 genes. (Supplementary Tables 19, 21). The program DATES yielded time estimates for the snow sheep and argali introgression events of 3,481 and 2,493 generations ago, respectively. Similar estimates were obtained with Ancestry_hmm: 3,096 and 2,545 generations ago (Supplementary Fig. 16). With a generation time of 4 years for Asiatic mouflon, both methods indicated that the introgression from snow sheep, as well as of bighorn and thinhorn sheep, occurred before the domestication 13,924 – 11,580 years BP. In contrast, the introgression from argali to Asiatic mouflon at 9,972 – 10,180 years BP coincides with the domestication process. Because introgression of argali in domestic sheep is confined to sympatric populations ^16,38^, we believe that the gene flow between argali and Asiatic mouflon did not take place until after the domestication process, resulting in the first domestic sheep lacking gene flow from argali. The gene flow from argali was probably also absent in the mouflon population ancestral to domestic sheep. GO categories and KEGG pathway of snow sheep and argali introgression into Asiatic mouflon were reported in the Supplementary Table 20.

**Figure 5.**
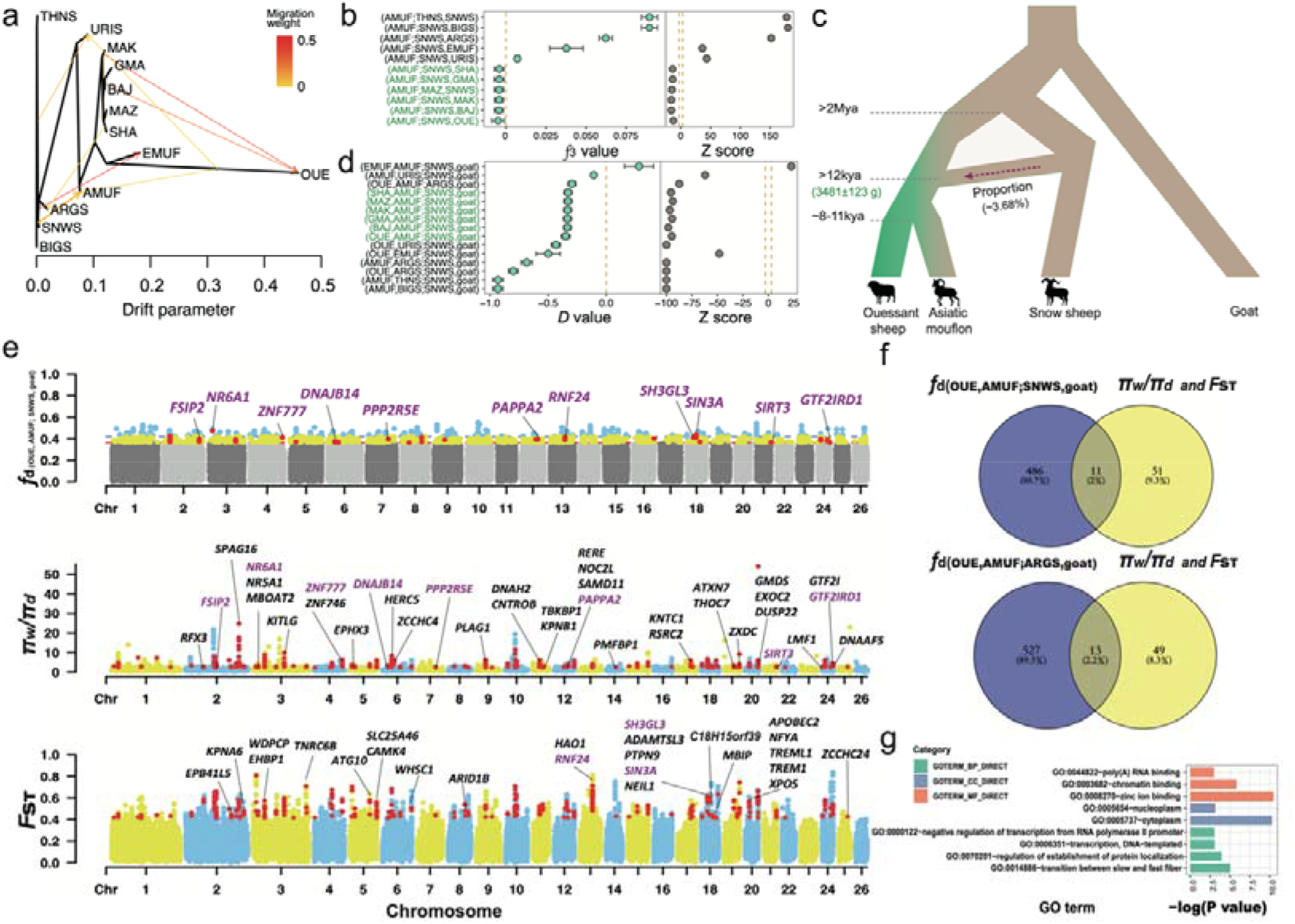
Local inference and annotation of introgression signals from snow sheep to Asiatic mouflon. **(a)** Treemix analysis when m=9. **(b, d)** *f*3 statistics and *D* statistics of Asiatic mouflon (AMUF) with snow sheep (SNWS) pairs. Double dashed line marked as the range of threshold from −3 to 3. **(c)** Demographic diagram of admixture from snow sheep (*O.nivicola*) to Asiatic mouflon (*O.orientalis*). **(e)** Introgressed regions identified in the Asiatic mouflon genome. A modified *f*-statistic (*f*_d_) for (OUE,AMUF;SNWS,goat), π ratio (π_w_/π_d_, i.e. π of Asiatic mouflon/and π of all the domestic sheep) and *F*_ST_ between Asiatic mouflon and all the domestic sheep for 100-kb windows with 20-kb steps is plotted along the chromosomes. Each dot represents a 100-kb window. For *f*-statistic, green and blue dots above the red horizontal line correspond to the FDR 5% and FDR 1% significance level thresholds, respectively. The regions containing genes among three indexes (*f*_d_, π ratio and *F*_ST_) are plotted in red dots. For the π ratio (π_w_/π_d_) and *F*_ST_, 340 domestic selection related windows are plotted, and 62 verified candidate domestication genes are marked in the plot. Overlapped genes (n=11) with *f*-statistic are marked in purple. **(f)** Venn diagram of overlapping genes (n=11 for snow sheep and n=13 for argali) between introgressed genes (n=497 for snow sheep and n=540 for argali) and candidate domestication genes (n=62). **(g)** GO enrichment for 62 overlapped domestication genes with the previous studies.

### Selection signatures in domestic sheep

To detect selection signatures in domestic sheep, we used pairwise differences π ratio (π_w_/π_d_) > 2.36 and *F*_ST_ between domestic sheep and Asiatic mouflon. We selected the overlap of the top 10% outliers in both methods, identifying 340 windows as candidate regions for selection. These regions contained a set of 131 selective functional genes (Supplementary Table 22 and Figs. 5e, f) which were significantly (*P* □ 0.05) enriched for GO terms involved in the activation of the innate immune response, positive regulation of defense response to virus by host, ectoderm development, membrane transport and enzyme activity (Fig. 5g).Of the 131 genes, 62 (47%) overlapped with domestication-related genes of previous studies, and were defined as the candidate domestication genes in sheep (Supplementary Table 23). Remarkably, from these candidate domestication genes 11 and 13 (15,365 genes on autosomes, significantly overlapped between two gene lists with Fisher’s exact test, *P* < 0.01) have been introgressed into Asiatic mouflon from snow sheep and argali (Figs. 5e, f). These genes were functionally involved in immune response (*HERC3* and *NFYA*), visual evolution (e.g., *RNF24*), resistance to virus (e.g., *SIN3A*) ^39–42^, production and reproductive traits [e.g. milk and protein yield (*SH3GL3* and *PAPPA2*)], fecundity (*DNAJB14* and *FSIP2*), body measurement (*SIRT3* and *SH3GL3*), tail type (*HAO1*), regulation of osteogenesis (*GTF2I*), skeletal muscle development (*ZNF777*) and lumbar vertebrae number traits (*NR6A1*) ^43–52^, and environmental adaptation [e.g., superior heat tolerance (*PPP2R5E*, *GTF2IRD1* and *DNAJB14*)] ^53–55^.

Of special interest was the introgressed genomic region chr3: 10980301-11211252 which contains gene *NR6A1* and had the highest (OUE, AMUF; SNWS, goat) *f*_d_ value (Fig. 6). We also computed the mean pairwise sequence divergence (*d_xy_*) of snow sheep and Asiatic mouflon or Ouessant sheep. This region also had a reduced mean pairwise sequence divergences (*d_xy_*) of snow sheep and Asiatic mouflon, a high *d_xy_* of snow and Ouessant sheep, and a low differentiation (*F*_ST_) of Asiatic mouflon and snow sheep, all indicating introgression of snow sheep into Asiatic mouflon (Figs. 6a – c).

**Figure 6.**
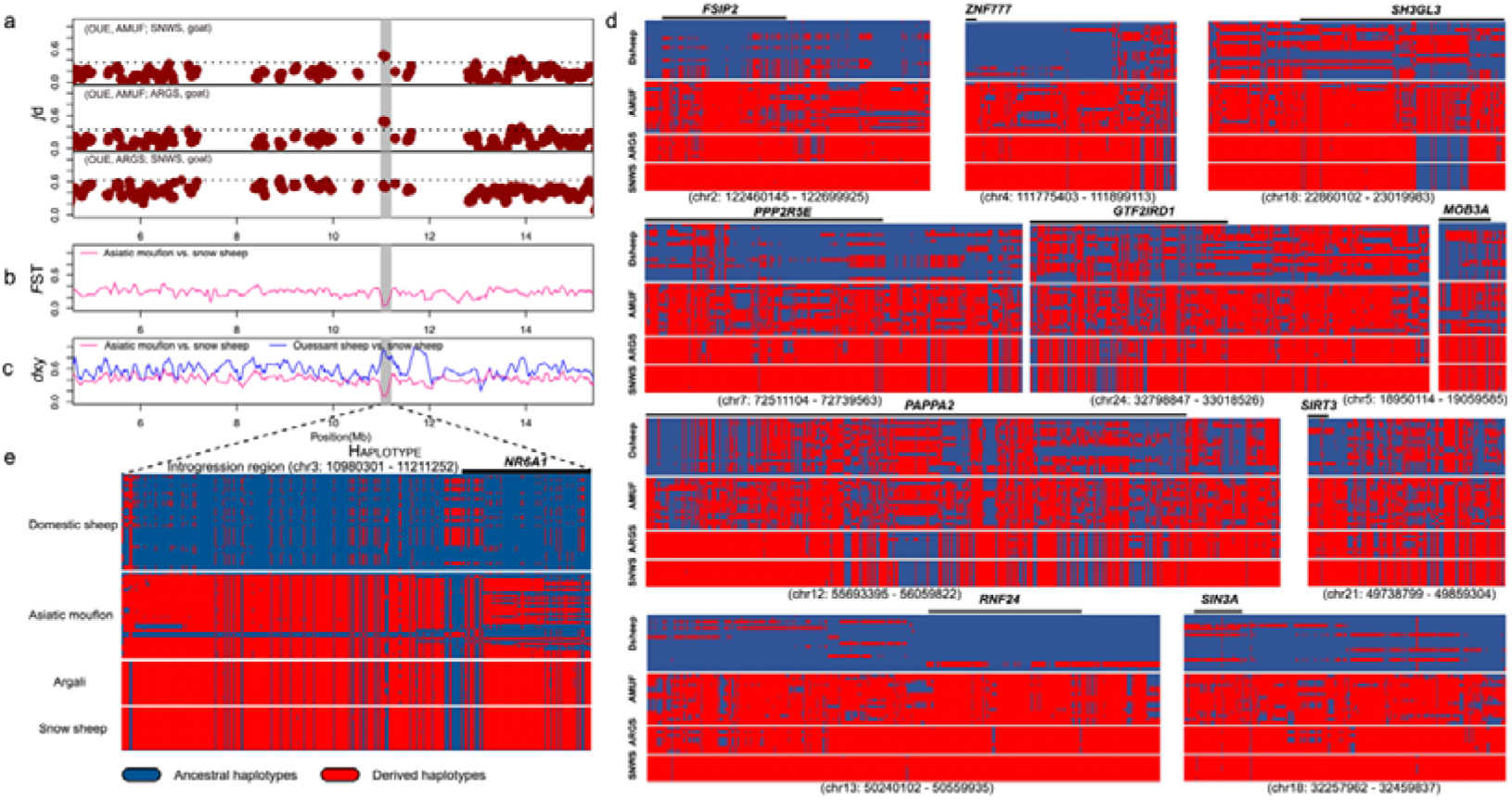
Local inference of genomic region at genes introgressed from snow sheep (SNWS) into Asiatic mouflon (AMUF). **(a)** *f* statistics (*f*_d_) based on (OUE, AMUF; SNWS, goat) comparison with (OUE, AMUF; ARGS, goat) and (OUE, ARGS; SNWS, goat) calculated for 100-kb windows with 20-kp steps across the genome for Asiatic mouflon. Each dot represented a 100-kb window, and the dashed line indicated the significance threshold (*P* < 0.05). **(b)** Population differentiation (*F*_ST_) around the introgressive genomic region between recipient (Asiatic mouflon) and donor (snow sheep). **(c)** Mean pairwise sequence divergence (*d_xy_*) of the introgression region between snow sheep and either Asiatic mouflon or Ouessant (OUE) domestic sheep population. **(d, e)** Haplotype patterns among all the domestic sheep, Asiatic mouflon, argali and snow sheep for the 11 genomic regions. Genes within the introgressed segments were marked upon haplotypes.

For the 11 genes introgressed from snow sheep into Asiatic mouflon, comparisons of haplotypes of the SNPs in the introgressive regions from argali, snow sheep, Asiatic mouflon and domestic sheep were shown in Figs. 6d, e. Notably, we found that the haplotype patterns of Asiatic mouflon strongly resembled those of snow sheep and argali, but differed strikingly from the patterns observed in the domestic sheep (Figs. 6d, e). Haplotype patterns showed most of the introgressive haplotypes of genes (e.g., *NR6A1, FSIP2, ZNF777, RNF24, PPP2R5E*) have not been selected and fixed in domestic sheep (Figs. 6d, e). Since most of the domestication-related genes are associated with production traits, this scenario could be explained by that introgressions associated with adaptation rather than production traits have been mostly selected in the genetic improvement stage after domestication ^38^.

### Common introgressions between wild and domestic sheep

In the introgression test (*D*-statistics and TreeMix analysis) between wild and domestic sheep, we found significant signatures of gene flow from (i) European mouflon into Ouessant sheep (OUE), (ii) urial and Asiatic mouflon into Shal (SHA), and (iii) argali into Tibetan sheep (GMA) (Fig. 4a, Supplementary Table 17, Supplementary Figs. 13, 17). We detected 154 genes located in the introgressed tracts from European mouflon to Ouessant sheep. These genes were significantly enriched in the GO terms and KEGG pathways of nerve conduction and development (*e.g.*,GO:0007269, GO:0098793, GO:0043065, GO:0090129, GO:0051965 and oas04360), cell adhesion (GO:0007156), intracellular signal transduction (GO:0035556 and oas04024) and walking behavior (GO:0007628) (Supplementary Table 24).

We identified regions containing 516 and 430 introgressed genes from the Asiatic mouflon or urial into Shal sheep, 251 of these were shared between the two species (Supplementary Table 24 and Supplementary Fig. 17). All these genes were significantly (*P* < 0.05) enriched in the GO terms with functions in tissue and organ development, reproduction and morphological change. In these tests of introgression from wild sheep to their sympatric domestic relatives, shared signals were detected in 10 functional genes (e.g., *CCDC67*, *FAT3*, *PCDH15* and *NEURL1*). These 10 common genes have functions associated with arid environment adaptation (*FAT3*) ^56^, immune response (*PCDH15*) ^57^, nervous response (*NEURL1*) ^58^ and disease susceptibility like noise-induced hearing loss (*PCDH15*) ^59^. Moreover, the genes introgressed from argali to Tibetan sheep were significantly enriched for GO terms in olfactory bulb development (e.g., *AGTPBP1*, *CRTAC1* and *RPGRIP1L*) and synaptic transmission (e.g., *GRIK2*, *PARK2* and *SHC3*) (Supplementary Table 24). Notably, two introgressed genes from Asiatic mouflon to Shal sheep (i.e., *RFX3* and *DNAJB14*) and one introgressed gene from argali to Tibetan sheep (i.e., *CAMK4*) were identified to be under domestication (Supplementary Tables 22, 23 and 24).

### Probability of incomplete lineage sorting (ILS)

We estimated the probability of incomplete lineage sorting (ILS) for the introgressed tracts identified from argali and snow sheep into Asiatic mouflon. The expected length of a shared ancestral tract (see Online methods) is *L*_snow_= 1/(1.5 × 10^−8^ × (2.3 × 10^6^ × 2) /4) = 57.97 bp, *L*_argali_= 1/(1.5 × 10^−8^ × (1.72 × 10^6^ × 2) /4) = 77.52 bp and the probability of a length of at least 96,410 bp and 98,037 bp (*i.e.,* the observed introgressed regions containing the domestication-related genes were 96,410 – 319,834 bp and 98,037 – 639,187 bp) is negligible (1 – GammaCDF (96,410, shape = 2, rate = 1/*L*) = 0) (Supplementary Table 25). Similarly, the probability of introgressed tracts appearing due to ILS detected from snow sheep to European mouflon and urial, as well as European mouflon, Asiatic mouflon and urial to domestic sheep were all approach zero (Supplementary Table 26). Thus, the inter-species introgressions detected above were unlikely due to ILS.

## Discussion

In this study, we generated a novel genomic dataset of high-depth whole-genome sequences of domestic sheep and all their wild relatives, including the wild ancestor of sheep (Asiatic mouflon), and the vulnerable (e.g., urial) and near threatened species (argali and Asiatic mouflon) according to the International Union for Conservation of Nature (ICUN) Red List. Our genomic data included different types of molecular markers such as SNPs, SVs, CNVs and INDELs, providing an important resource for the genetic improvement of sheep, as well as for ecological and evolutionary studies of the wild species.

This is the first comprehensive and in-depth investigation on phylogeny and introgressions among the whole *Ovis* genus. Different from previous studies ^1,60^, multiple up-to-date analyses were applied to cross-validate the obtained results. For example, to better understand the trajectories of connections between admixture events and phylogenetic relationship across the whole genome, we used sliding window-based and fitting-based methods to construct the consensus trees. Additionally, we implemented the introgression tests based on several statistical approaches such as *D*-statistics, *f*-statistics, TreeMix and admixture analyses. Further, we verified the introgression events (including introgression sources and time) using the admixturegraph fitting method and dated the introgression time using both model-based and LD-decay based methods. All the analyses showed accordant results.

We verified our SNPs based on both statistical and experimental methods, securing the dataset for the subsequent analysis. By comparing with other species, we found on average 17.61 million SNPs/per individual (9 – 21 SNPs/kb among the *Ovis* species), which is less than that in goat (53 – 54 SNPs/kb) ^61^, but higher than that in swine (1 SNP per 10.3 kb) ^62^.

Within the *Ovis* species, a relatively low diversity and effective population size (4,000 – 10,000) in the Pachyceriforms may be ascribed to long-term geographic and genetic isolation ^1,21^ and is relevant for its conservation. The much less genetic diversity observed in Pachyceriforms than that in Moufloniforms could be due to (*i*) the common ancestor of Pachyceriforms should have migrated out from Eurasia, the distribution region of Moufloniforms, with genetic drift and differentiation between each other; and (*ii*) the genome of domestic sheep has been used as the reference for SNP mapping, while domestic sheep is phylogenetically further from Pachyceriforms than Moufloniforms. We observed the highest diversity in Asiatic mouflon (π = 0.0044), which was much higher than in domestic sheep (π = 0.0032). This has been observed previously ^63^ and can be explained by the domestication bottleneck ^64^. Our results also showed lower diversity estimates than previous investigations using whole-genome BeadChip SNPs and mtDNA variation^63,65^. Higher estimates from the BeadChip could be explained by the ascertainment bias in the chip design.

Previous molecular evidence for taxonomic classification has so far mostly been based on mtDNA sequences ^1,60,66^. However, pervasive and frequent autosomal introgressions ^14,16^ probably accounts for the lower estimates of the coalescence time compared with those from mtDNA sequences ^1,60^. In particular, the evidence from SMC++, CoalHMM statistics, phylogenetic trees, admixture analysis, and mean population differentiation (*F*_ST_) index indicated a more recent divergence of European mouflon and domestic sheep (∼5,000 BP), than estimated on the basis of mtDNA sequences (∼21,000 BP) ^67^. The recent divergence of European mouflon from domestic sheep was also supported by archaeological data ^24^. Our evidence confirmed that European mouflon emerged as feral domestic sheep when the earliest wave of domestic hair sheep, was displaced by a second wave of wool sheep ^15,68,69^.

Also, we obtained a more recent split time between argali and domestic sheep (∼0.12 - 0.15 Mya) than earlier estimates. The earlier divergence time was based on orthologous genes using PAML and the node was calibrated using four fossil records such as the divergence of the opossum and human (124.6–134.8 million years ago [Mya]), human and taurine cattle (95.3–113 Mya), taurine cattle and pig (48.3–53.5 Mya), and taurine cattle and goat (18.3–28.5 Mya) ^70^. Also, different estimates of 1.72 ± 0.36 Mya and ∼ 2.93 Mya were obtained from mitochondrial sequence variations ^1,60^ using the five fossil calibration time of 18.3–28.5 Ma between Bovinae and Caprinae, 52–58Ma between Cetacea and hippopotamus, 4 34.1 Ma between baleen and toothed whales, 42.8–63.8 Ma between Caniformia and Feliformia, and 62.3–71.2 Ma between Carnivora and Perissodactyla. This difference could be due to different mutation rates of the whole-genomes and mtDNA sequences, and different calibration time points have been used in different studies. Additionally, we used two model of coalHMM (Isolation with migration and Isolation model) with full consideration of migration after speciation, and the estimates have always been lower than those estimated by mtDNA sequences and protein-coding genes ^71^. Besides these, the more recent divergence time estimated here could be attributed to the extensive genomic introgressions between the sequenced genomes of the two species.

Remarkably, the Admixture and Treemix patterns, as well as *D*-statistics and *f*_d_ statistics consistently showed introgression of the Pachyceriforms, comprised by the snow sheep and its American relatives (bighorn and thinhorn sheep), into European mouflon. The Pachyceriforms also introgressed Asiatic mouflon, but this was a more recent event and involved a different set of genomic segments (Supplementary Fig. 15). The introgression percentage as inferred by *f*_d_ statistics were 1.47%, 1.45% and 1.43% from bighorn, thinhorn and snow sheep to European mouflon were, while higher introgression percentages of 3.7%, 3.63% and 3.68% were from bighorn, thinhorn and snow sheep to Asiatic mouflon. This indicated that the wild ancestors of the European mouflon, and consequently also the first hair sheep domesticated, descend from a population that differs from the Asiatic mouflons in this study, which were sampled in Iran. In contrast, the Iranian Asiatic mouflons are phylogenetically diverse and close to urial, which is in line with the mtDNA (Supplementary Fig. 7) and Y-chromosomal phylogeny ^19^. As the range of snow sheep and their American relatives did not extend to Europe, our results suggested that European mouflons may have partially descended from a now extinct sheep in Europe and arose through hybridization events between this species and feral domesticated sheep (Fig. 4, Supplementary Fig. 7). Some wild sheep species live in extreme environments, such as snow sheep in the extreme cold arctic regions, and argali on the cold Qinghai-Tibetan Plateau and the Pamir highland. Thus, our data may be relevant for environmental adaptation. However, it’s challenging to confirm the ancient introgression trajectories based on modern samples, ancient samples of *Ovis* species are demanding to answer this question.

Recently, there has been a strong interest in inter-species introgression, particularly from wild relatives to domestic animals such as pig, goat and sheep ^9,10,13,15^. For example, an earlier study has shown adaptive introgression and selection on domestic genes in goat ^72^. In particular, *MUC6* was found to be introgressed from a West Caucasian tur-like species into modern goat during domestication, and is nearly fixed in domestic goat with the function of pathogen resistance ^72^. In the *Ovis* genus, hybridization among species has been documented in previous field and molecular studies ^6,9,66^. However, adaptive introgression from distantly related wild species into the wild ancestors of domestic animals or into domestic animals has rarely been investigated ^72,73^. Genomic signature of adaptive introgression from European mouflon into domestic sheep has been previously reported ^15^. An earlier whole-genome SNP analysis suggested that historical introgression from wild relatives is associated with climatic adaptation and that introgressed alleles in *PADI2* have contributed to resistance to pneumonia in sheep ^38^.

A strong signature of adaptive introgression from argali into Tibetan sheep was detected, and the introgressive genes involved in hypoxia and ultraviolet signaling pathways (e.g., *HBB* and *MITF*) and associated with morphological traits such as horn size and shape (e.g., *RXFP2*). The introgressed genes were related to adaptation to the extreme environment in the Qinghai-Tibetan Plateau ^16^. We also identified other genes in Tibetan sheep introgressed from argali, associated with disease resistance to pathogens (e.g., *ACTN4*), and with olfactory development (*e.g.*, *AGTPBP1*), and locomotion (*e.g.*, *OXR1*), possibly related to adaptation to the semi-wild grazing and anoxic environments in plateau. Furthermore, we found patterns compatible with adaptive introgression from the Pachyceriform sheep and argali into urial, Asiatic mouflon and European mouflon. However, it is challenging to validate the function of these genes the in vivo or vitro in the wild animals.

We detected adaptive introgression from various wild species into Asiatic mouflon, covering several domestication-related genes. Inspection of these domestication-related genes (e.g., *KITLG*, *CAMK4*, *NR6A1*, *RNF24*, *MBIP*, *SH3GL3*, *GMDS*, *EXOC2* and *GTF2I*) indicated their functions associated with important morphological, physiological and production traits such as litter size and mammary cycle ^74,75^, early body weight (e.g., *PLAG1* ^76^), regulation of follicular development (e.g., *NR5A1*; ^77^) in sheep. Theoretically, particular functions of these domestication-related candidate genes indicated relevant traits have been the targets under intensive selective pressure during the domestication process, which eventually led to emergence of the typical morphological, production, physiological and behavioral differences between domestic sheep and their wild ancestors ^78^. In practice, the highly differentiated nonsynonymous mutations in coding regions of the genes should be functionally important and could be integrated in marker-associated selection and genomic selection for related traits in future genetic improvement of domestic sheep^17^.

## Conclusions

In conclusion, we estimated the phylogenetic relationships of the sheep species on the basis of high-depth whole genome sequences. Our results suggested a feral origin of domestic sheep for European mouflon around 6,000 – 5,000 years BP and a genetic overlap of urial and the Iranian Asiatic mouflon. We found extensive introgression events among the *Ovis* species, which partially overlap with regions under selection in domestic sheep. Our results provide novel insights into changes in the genome landscapes of domestic sheep and their wild ancestor occurring during and after domestication.

## Online Methods

### Samples and DNA extraction

Seventy-two whole-genome sequences of 6 domestic breeds (*n* = 18) and all wild species (*n* = 54) of the genus *Ovis* were included in this study (Supplementary Table 1). Here we followed the classification of Nadler et al. (1973) due to the greatest taxonomy number. The classification was based on morphological traits in chromosome diploid number. These domestic breeds were selected from sheep which have showed genomic introgressions from sympatric wild relatives ^15,16,38^. Thirty-five whole-genome sequences were sequenced in this study and 37 were from our previous studies ^17,19^. The 35 genomes generated here consisted of 7 domestic sheep from 3 populations, including Tibetan sheep (GMA, Maqu county, Gansu), Mazekh sheep (MAZ) in Azerbaijan and Makui sheep (MAK) in Iran, and 28 wild sheep from 5 species, including *O. musimon* (*n* = 3), *O. vignei* (*n* = 5), *O. nivicola* (*n* = 8), *O. dalli* (*n* = 6) and *O. canadensis* (*n* = 6), all of which were understudied in previous studies. The 37 public genomes comprised 11 domestic sheep from the following breeds: 3 French Ouessant (OUE) sheep, sampled in the Netherlands, 3 Baidarak sheep (BAJ) from Russia, 3 Shal sheep (SHA) from Iran, 1 Tibetan sheep and 1 Makui sheep, as well as 26 wild sheep genomes from 3 species (*O. orientalis*, *n* = 16; *O. ammon*, *n* = 8 and *O*. *vignei*, *n* = 2; Supplementary Table 1). Historical information, geographic distribution, and morphological traits such as body size, horn morphology, color and pattern of the coat have been used in the definition of species ^79^ and types and varieties of hair and wool sheep ^27^. Genomic DNA was extracted from the blood or tissue samples using the standard methods of proteinase *K* solution and phenol-chloroform extraction ^80^. DNA samples with a clear band in sepharose gel, an OD_260_/OD_280_ ratio between 1.7 and 2.0 and a concentration at least 20 ng/ L were used for the library construction.

### DNA sequencing and read filtering

Whole-genome sequencing was performed using the Illumina Hiseq Xten. At least 1.5 μg of genomic DNA from each sample was sheared to a 180-500 bp range using the Covaris S220 instrument (Covaris, Woburn, MA, USA) and used for Illumina library preparation. Sequencing libraries were constructed using the Truseq Nano DNA HT Sample preparation Kit (Illumina Inc., San Diego, CA, USA) following the manufacturer’s instructions. In brief, DNA fragments were end-repaired, A-tailed, ligated to paired-end adapter, and the fragments with ∼350 bp insert length were selected for amplification by 8-12 cycles of PCR using the Platinum Pfx Taq Polymerase Kit (Invitrogen, Carlsbad, CA, USA). PCR products were purified with the AMPure XP system (Beckman Coulter, Brea, CA, USA), and libraries were analyzed for the size distribution by the Agilent 2100 Bioanalyzer (Agilent Technologies, Palo Alto, CA, USA) and quantified in real-time PCR. The constructed libraries were sequenced on the Illumina HiSeq X Ten platform (Illumina Inc.) and paired-end 150 bp reads were generated.

All the newly generated and retrieved whole genomes (*n* =72) were included in the following analyses. On average, 95.83% of the sheep reference genome was covered by the depth of ≥4×, 90.11% was covered by ≥10×, and 46.68% was covered by ≥20×.To obtain reliable reads, we removed the raw paired-reads that meet any of the following three criteria: (*i*) unidentified nucleotides (N-content) ≥ 10%; (*ii*) reads pair with adapters; and (*iii*) > 50% of the read bases with a phred quality (Q) score less than 5.

### Reads mapping, variant detection, quality control and annotation

Clean reads were mapped to the sheep reference genome OARv4.0 (GCA_000298735.2) using the BWA v0.7.17 MEM module ^81^ with the parameters bwa -k 32 -M -R. Duplicates were removed using Picard MarkDuplicates and sorted using Picard SortSam (https://broadinstitute.github.io/picard/). To obtain reliable alignments, the reads meeting any of the following three criteria were filtered: (*i*) unmapped reads; (*ii*) reads not mapped properly according to the aligner used above; and (*iii*) the reads with RMS (root mean square) mapping quality < 20. Base quality score recalibration (BQSR) with ApplyBQSR module (default parameters) was used to detect the systematic errors during the sequencing process.

Variant discovery was carried out using the Genome Analysis Toolkit (GATK-v4.0.4.0) best practices pipeline, followed by a joint genotyping method on all samples in the cohort ^82^. In summary, we firstly called the variants based on each sample using Haplotypecaller module in GVCF mode with the parameter - genotyping-mode DISCOVERY --min-base-quality-score 20 --output-mode EMIT_ALL _SITES --emit-ref-confidence GVCF. Then, we implemented the joint genotyping procedure by consolidating all the GVCFs with the GenotypeGVCFs module. Furthermore, we combined all the variants using CombineGVCFs. Variant sites were identified for each of the eight species, separately. Within each species, the following successive filtering processes were applied for the variant site and genotype quality control: First, raw SNPs were hard filtered using the VariantFiltration module with the strict parameters -filter-expression QUAL < 30.0 || QD < 2.0 || MQ < 40.0 || FS > 60.0 || SOR > 3.0 || HaplotypeScore > 13.0 || MQRankSum < −12.5 || ReadPosRankSum < −8.0 in each species, separately. We then merged all the 8 variant datasets from eight species using the bcftools merge function after the bcftools index. In addition, PLINK v1.9 ^83^ was used to filter SNPs which meet any of the following criteria: (*i*) proportion of missing genotypes among all the individuals over 10% (geno 0.1); (*ii*) SNPs with minor allele frequency (MAF) higher than 0.05 (maf 0.05); (*iii*) SNPs showing an excess of heterozygosity (--hwe 0.001); and (*iv*) non-biallelic sites. This yielded a high-quality set of variants including 6,558,545 SNPs were obtained for the genomic introgression and diversity analyses. For analyzing population genetic structure, we excluded SNPs in LD with *r*^2^ ≥ 0.2 (--indep-pairwise 50 5 0.2). All the SNPs were annotated using the ANNOVAR v.2013-06-21 software^84^ and phased using Shapeit v4.1.3 ^85^.

### SV detection and annotation

To identify reliable structural variants (SVs), we detected the SVs by implementing four independent calling pipelines. First, SVs were detected based on the filtered and sorted BAM file using novoBreak v.1.1.3 ^86^, which detects deletions (DEL), inversions (INV), tandem duplications (DUP) and inter-chromosomal translocations (TRA). Second, SVs were identified using configManta.py in manta v.1.6.0 ^87^. Manta reports SVs as deletions (DEL), inversions (INV), tandem duplications (DUP), insertions (INS) and inter chromosomal translocations (TRA). Third, SVs were detected using GRIDSS v2.6.2 ^88^. SV files in VCF format were then annotated using a custom R script (https://github.com/PapenfussLab/gridss/blob/master/example/simple-event-annotation.R). GRIDSS generates the same variant types of SVs as those by manta. These three pipelines utilized the same input of 72 BAM files. Fourth, paired-end reads were re-mapped to the sheep reference genome (Oar_v4.0) using the align module of SpeedSeq v.0.1.2 ^89^.

In addition, sorted and duplicate-marked BAMs, which contain split reads and discordant read-pairs, were generated. SVs were then identified from the split reads and discordant pairs using LUMPY v.0.2.13 ^90^. CNVs were detected from the difference in read depth using CNVnator v.0.3.3 ^91^. The inferred breakpoints by LUMPY were genotyped using SVTyper v.0.1.4 ^89^. The variant types of SVs detected by the SpeedSeq framework are the same as those by the GRIDSS pipeline. In these two pipelines, we generated non-uniquely mappable genomic regions for autosomes and X chromosomes, respectively, using SNPable (http://lh3lh3.users.sourceforge.net/snpable.shtml), and these regions were masked in the SV detection by the two methods described above.

To reduce the false positive rate, SVs in both autosomes and X chromosomes from the four strategies (novoBreak, manta, GRIDSS and SpeedSeq) which meet the following seven criteria were retained: (*i*) at least three split reads (SR) or three spanning paired-end reads (PE) supporting the given SV event across all the samples; (*ii*) SVs with precise breakpoints by novoBreak (flag PRECISE); (*iii*) SVs passing the quality filters suggested by NovoBreak, manta and GRIDSS (flag PASS); (*iv*) SVs with more than four supporting reads (flag SU) and without ambiguous breakpoints (flag IMPRECISE) in SpeedSeq; (*v*) SVs with lengths between 50 bp and 1 Mb; (*vi*) SVs without intersections between different variant types; and (*vii*) SVs identified by at least two pipelines. For each sample, the shared SVs detected at least by two of the four independent pipelines were merged using SURVIVOR v.1.0.6 ^92^ with the parameters 500 2 1 1 0 50.

SVs were annotated based on their start positions using the package ANNOVAR v.2013-06-21 ^84^. Species-unbalanced SVs are defined as SVs which are unevenly distributed among different species. A two-sided Fisher’s exact test was utilized to determine whether the distribution of each SV is uniform. The *P*-values for all the SVs were calculated with the Fisher.test function in R followed by the Benjamini– Hochberg false discovery rate (FDR) adjustment. SVs with FDR < 0.05 were considered as species-unbalanced.

### SNPs and CNVs validation

74 randomly selected SNPs of 4-12 individuals were verified by PCR amplifications and Sanger sequencing. The primers used for the PCRs were designed with the software Primer Premier 5 ^93^. The PCR reactions were performed in a total volume of 25 μl, consisting of 12.5 μl 2× Taq MasterMix (Kangwei, Beijing, China), 2 μl (10 pmol/µL) reverse and forward primers, 1 μl template DNA (30 ng/µL) and 9.5 μl double-distilled water (ddH_2_O) under the reacting condition of initial denaturation at 95 °C for 3 min, 35 cycles for the following three steps, such as denaturation at 95 °C for 15 sec, annealing at 60 °C for 15 sec, and extension at 72 °C for 30 sec, with a final extension at 72 °C for 5 min. Following the PCR, the amplification products were sequenced on the Applied Biosystems 3730XL DNA Analyzer (Life Technologies, Carlsbad, CA, USA), and the sequencing peaks were checked with the software SEQMAN module of DNASTAR’s LASERGENE ^94^. Subsequently, genotypes obtained from the Sanger sequencing were compared with those inferred by the GATK pipelines (described above) from resequencing data for the same individuals.

Moreover, 14 randomly selected CNVs (e.g., seven deletions and seven duplications; Supplementary Table 13) were validated by quantitative real-time PCR (qPCR) or PCR. Primers designed surrounding the deletions and within the duplications with the software Primer Premier 5 (Supplementary Table 13). Deletions were genotyped by PCR amplification and agarose gel electrophoresis. We measured the relative copy numbers of one deletion and all duplications using qPCR on the QuantStudio^TM^ 6 Flex Real-Time PCR System (Life Technologies, Carlsbad, CA, USA) using SYBR Green kit (Promega, Madison, WI, USA). Following a previous study on sheep ^95^, *DGAT2* gene was used as the internal reference gene. qPCR reaction was in 25 μl volume consisting of 12.5 μl 2× SYBR Green qPCR Mix (Life Technologies, Carlsbad, CA, USA), 1 μl (10 pmol/µL) each primer (forward and reverse), 2 μl template DNA (30 ng/μl), and 8.5 μl ddH_2_O. The thermocycling condition includes an initial denaturation at 95 °C for 10 min, 40 cycles for the next three steps, such as denaturation at 95 °C for 15 s, annealing at 60 °C for 15 s and extension at 72 °C for 1 min, and a final extension at 72 °C for 10min.

For qPCR, the ΔΔ*C*_T_ method ^96^ was applied to estimate the relative copy numbers. Equation for ΔΔ*C*_T_ value is ΔΔ*C*_T_ = [(*C*_T segment_ □ *C*_T_*DGAT2*_)_target sample_ □ (*C*_T segment_ □ *C*_T_*DGAT2*_)_control sample_], where *C*_T segment_ is threshold cycle (*C*_T_) of target CNV segment and *C*_T_*DGAT2*_ is the *C*_T_ of the internal reference gene ^96^. We also measured the standard deviation of the ΔΔ*C*_T_ value using the formula: s = (s_1_^2^ + s_2_^2^)^1/2^, where s_1_ is the variance of target *C*_T_ value (3 replications) and s_2_ is the variance of the reference *C*_T_ value (3 replications). The value of 2×2^-ΔΔ^*^C^*^T^ between 1.5 and 3 were considered to most likely represent a normal copy number of 2, below 1.5 or above 3 are considered as deletions or duplications, respectively ^20^. This was used to evaluate the concordance of calling results obtained from four SVs calling strategies and the relative copy number from the qPCR.

### Inference of demographic history

We inferred past temporal change in *Ne* and population split times using the pairwise sequentially Markovian coalescent (PSMC) modelling (http://github.com/lh3/psmc) and SMC++ program (https://github.com/popgenmethods/smcpp#masking). We applied the parameters of a generation time (*g*) of 3 years, neutral mutation rate (μ) = 2.5×10^−8^ per base pair per generation, a per-site filter of ≥ 10 reads and no more than 25% of missing data ^97^, including only autosomes from one high-coverage genomes (> 18×) per species (PSMC, Supplementary Table 2) or three individuals per population or species (SMC++). We performed 1,000 bootstrapping simulations to estimate the variance of *Ne*.

### Genomic diversity and population differentiation

For each individual, genome-wide nucleotide diversity was calculated based on the set of high-quality SNPs (*n* = 6,558,545) using Vcftools v0.1.13 with a window size of 200-kb. Genome-wide pairwise *F*_ST_ and *d*_xy_ genetic distance matrices between populations was estimated using in-house python scripts with a window size of 100-kb and a 20-kb step size. The matrices of pairwise distances were then plotted using the Corrplot package of R. In order to assess the genome-wide LD patterns of each species, we calculated *r*^2^ value using the program PopLDdecay v3.30 ^98^ (https://github.com/BGI-shenzhen/PopLDdecay) with the default parameters and after filtering the sites with more than 10% missing genotypes among the individuals of each species cohort.

### Population genetic structure and phylogenetic reconstruction

We implemented principal components analysis (PCA) using the Smartpca program ^99^ in the software EIGENSOFT v7.2.1 ^100^ without outlier removal iteration (numoutlieriter: 0) but with the default settings of the other options. The Tracy-Widom test was used to determine significance of the eigenvectors. The first two eigenvectors were plotted. We used the Ohana tool suite ^101^ to infer the global ancestry and the covariance structure of allele frequencies among the species. The number of ancestry components (*K*) was set in a range from 2 to 11. For each *K*, we terminated the iteration when the likelihood improvement is smaller than 0.001 (-e 0.001). We only reported the ones which reached the best likelihood for each *K*. Population trees at each *K* (Supplementary Fig. 12b) were plotted using the program Nemetree (http://www.jade-cheng.com/trees/).

The phylogenetic tree of the nine species was constructed using the maximum likelihood method implemented in the RAxML v8.2.3 ^102^ with the multiple nucleotide substitution models. The tree was inferred based on the 12,837 protein coding sequences (CDS) on autosomes and 513 CDS on X chromosome, separately. We used the protein-coding gene annotation file from NCBI (ftp://ftp.ncbi.nlm.nih.gov/genomes/Ovis_aries/GFF/). Only CDS with length multiple of 3 were considered in the phylogenetic inference. The consensus trees (Supplementary Fig. 5) based on the whole genome was built on the concatenated CDSs of autosomes (33,868,497 bp), X chromosome (1,331,184 bp) and the whole mitogenomes (16,616 bp), respectively. Moreover, seventy-two haploidized whole-genome sequences for all the individuals were generated using the -doFasta3 option in ANGSD ^103^ (Fig. 2b and Supplementary Fig. 7), which uses the bases with the highest effective depth (EBD) and considers both mapping quality and scores for the bases ^104^. To examine the impact of different assembly methods on the phylogenetic inference, we also tested the options of -doFasta 1 and -doFasta 2, which utilize the genomic sites by randomly selecting the base or selecting the base with the highest depth.

The preliminary tree for the optimization were constructed using the GTRCAT model in RAxML. Phylogenetic inference of autosomal and X chromosomal sequences was then implemented based on the first two codon positions and the third codon position of the whole concatenated coding sequence using the GTRGMMA model in RAxML. The final trees after 200 bootstrapping replicates were generated using GTRCAT model in RAxML and returned to the preliminary tree labeled with bootstrap values. To clarify discordant coalescent events among different tracts in the genome, we split the whole genome into 1-Mb tracts, which result in 2,598 non-overlapping windows, respectively. We inferred the ML trees using the GTRGAMMA model. Finally, trees were built of each 1-Mb windows (Supplementary Fig. 6). Numeration (classification and ranking) of trees was conducted using all.equal function in R package of Ape (analyses of phylogenetics and evolution) and plotted by in-house R scripts. The trees of each tract of autosomes and X chromosome in 1-Mb window were fitted and visualized by Densitree v2.0.1 ^105^ (Fig. 2a). Mitochondrial sequences between sheep and goat were blasted using MEGA7 ^106^. Genomic coordinates of goat were transferred based on locations of the sheep genome after trimming the poorly mapped sites. Finally, we merged the 73 mitochondrial sequences in the phylogenetic analysis with goat as the outgroup.

### Estimation of split time

Divergence time was estimated locally based on each 1-Mb tracts across autosomes. We used the coalescent hidden Markov model (CoalHMM) ^23^, a framework for demographic inference using a sequential Markov coalescent method, to estimate the split time with or without migrations among species. We first converted the pairwise sequence alignments using python scripts prepare-alignments.py (https://github.com/birc-aeh/coalhmm/tree/master/scripts/). The I-CoalHMM and IM-CoalHMM models were then applied to the dataset of 1-Mb tracts. The two models utilized the genome alignments of two species to calculate the time of speciation. In the I-CoalHMM model, a prior of split time and ancestral effective population size were needed, whereas in IM-CoalHMM model extra migration rates were also needed. The recombination rate was set as 1.5 cM/Mb ^107^. We combined the pairwise alignments between species totaling nine pairs and used 1-Mb splitting windows of the whole genomes for each pair and discarded the windows with > 10% missing bases. We filtered the time estimates for the windows using the following criteria: (*i*) a split time of below 1,000 years or above 10,000 years for European mouflon and sheep, below 1,000 years or above 20,000 years for Asiatic mouflon and sheep or below 10,000 years or above 10 million years for the divergence between the other seven pairs of species (Supplementary Figs. 9 – 11), (*ii*) a recombination rate below 0.1 cM/Mb or above 5 cM/Mb, and (*iii*) an ancestral effective population size below 5,000 or above 1,000,000 ^14^.

### Migration events by TreeMix analysis

To infer migration events among the eight species, we used TreeMix v1.13 to construct a ML tree with bighorn as the root using the “-noss” option to turn off the sample size correction, a window size (-*K*) of 500 SNPs (around 609-kb in this study) to account for the impact of LD, which is more than the average LD length of approximately ∼150-kb observed in sheep ^7^. Blocks with 500 SNPs were resampled and 100 bootstrap replications were performed. We constructed the ML trees with 0-11 migration events and corresponding residuals. The proportions of explained variance (Supplementary Fig. 14) for the migration numbers were calculated using in-house scripts ^108^.

### Gene flow among species

To infer the ancestral alleles, genomic comparison between domestic sheep (*O. aries*) and domestic goat (*C. hircus*) was carried out using the LAST v984 program (http://last.cbrc.jp/) (Supplementary Fig. 18). We aligned the sheep reference genome (Oar_v4.0) to the goat reference genome (ASR.1) while masking the repeat regions. Only autosomal one-to-one orthologs were considered in the alignment between the two species using the lastal module with the parameters of -m 100 –E 0.05. To visualize the corresponding orthologs between species, a synteny plot was created using the circlize function in the R package. Samtools mpileup and Bcftools call were then used to call ancestral alleles. We merged these ancestral variants with the combined SNPs of all the 72 samples using Bcftools merge after indexing the two datasets. The combined dataset was used to detect introgression among species.

To detect the potential gene flow among species, we conducted the ABBA-BABA test (*D-*statistics) based on two data panels: single high-depth genomes and high reliable SNPs among all the individuals. These two datasets can be collated between each other to reduce variants calling errors. For the first data panel, we performed the admixture analysis using ANGSD -doAbbababa 1 module with goat as the outgroup and the block size of 1,000,000 bp. For the second data panel, we examined the admixture among species using the qpDstats module of AdmixTools ^109^ and goat as the outgroup, which is a formal four-population test of admixture. Furthermore, we performed the three-population test using the qp3pop module of AdmixTools. The statistical significance of *D* value was evaluated using a two-tailed Z test, with |Z-score| > 3 to be significant ^110^. We built the admixture graphs, fitted the graph parameters and visualized the goodness of fit using admixturegraph ^111^ package in R.

### Inference of introgressed genomic regions

To further localize the introgressed genomic regions across the whole-genome, a window-based Patterson’s four-taxon *D*-statistic test *D* (P1, P2, P3, O) and modified *f*-statistic (*f*_d_) test with 100-kb length windows and 20-kb steps was performed using the methods of Martin *et al*. (2015) ^112^. P1 was the reference population with no gene flow with P3 and is closer to P2 than P3. Here, goat was used as the outgroup (O), which was the ancestral population and shared derived alleles with populations P1, P2, and P3. The significance level (*p*-value) of Z-transformed *f*_d_ value was corrected by multiple testing using the Benjamini–Hochberg FDR method ^28^. Windows with positive *D* values and *p* values (FDR adjusted) < 0.05 were selected as the significantly introgressed regions, and the adjacent windows were merged into concatenated introgressed regions ^113^.

We tested for genomic introgressions between different combinations of species.

(*i*) *D* (OUE, target; X, goat): The domestic population of Ouessant (OUE) serves as the reference population, the goat reference sequence was the outgroup, European mouflon, Asiatic mouflon or urial were the targets and X (bighorn, thinhorn, argali and snow sheep) was the to be tested source of introgression.

(*ii*) *D* (GMA, target; European mouflon, goat): We selected as reference the old and native Tibetan sheep (GMA) that has no potential gene flow with European mouflon and the targets were Ouessant in France, Mazekh in Azerbaijian, Makui and Shal sheep in Iran.

(*iii*) *D* (GMA, target; Asiatic mouflon, goat): the targets were the domestic Mazekh, Makui and Shal sheep.

(*iv*) *D* (OUE, Baidarak; snow sheep, goat): OUE from France was a suitable reference because its large distance to the range of snow sheep and Baidarak was the target.

(*v*) *D* (OUE, target; argali, goat): targets were domestic Tibetan sheep and Russian Baidarak.

(*vi*) *D* (OUE, target; urial goat): targets were domestic Mazekh, Makui and Shal sheep. In addition, we calculated mean pairwise sequence divergence (*d*_xy_) and *F*_ST_ value between the target population (P2) and the test population (P3), as well as between the test population (P3) and the reference population (P1). Introgression but not shared ancestry reduces *d_xy_* in the target regions ^112^. Similarly, introgressed regions have lower divergence (*F*_ST_) than other regions.

### Dating introgression events

We dated the time of ancient introgression using DATES ^114^ and Ancestry_hmm program ^115^. The software DATES computed the weighted LD statistic to infer the population admixture history, which has been developed for human datasets. However, because of the short generation time for sheep, the time estimates using DATES might be younger than expected. Thus, we also applied the Ancestry_hmm program ^115^, using phased data and only SNPs with at least two alleles in the reference populations and applying the following filters: (*i*) SNPs with allele frequency difference lower than 0.1 between the two reference populations; and (*ii*) SNPs with allele number less than 6 in a reference panel. Other parameters were set as default. We set the proportion of admixture (*m*) according to admixture fraction obtained above by the *f* statistics (*f*_d_) across the whole genomes. For dating the introgression from bighorn sheep, thinhorn sheep, snow sheep or argali as source of introgression (reference population 2) into Asiatic mouflon, we used urial as the ancestor (reference population 1). We applied a single pulse model for genotype data from each population and ran 100 bootstrap replicates using a block size of 5,000 SNPs.

### Incomplete lineage sorting

We calculated the probability of incomplete lineage sortings (ILSs) following the method in Huerta-Sánchez *et al*. (2014) ^116^. Briefly, the expected length of a shared ancestral sequence is *L*=1/(*r*×*t*). The probability of a length of at least *m* follows from 1 □ GammaCDF (*k*, shape = 2, *r* = 1/L), in which GammaCDF is the Gamma distribution function, *r* is the recombination rate per generation per bp, *m* is the length of introgressed tracts, and *t* is the length of the two species branch since divergence. According to the theoretical expectation, we can exclude the possibility of common ancestral source when the detected length of tracts (*m*) > *L* or the probability of a length of at least *m* infinitely approaches zero. Here, we set recombination rate of 1.5 × 10^−8 107^, generation time of 4 years for Asiatic mouflon and urial ^117^ and 3 years for domestic sheep ^118^. We set divergence times of 2.3 mya for snow sheep and Asiatic mouflon, 1.72 mya for argali and Asiatic mouflon ^21^, 2.42 mya for the Pachyceriforms and the Moufloniforms (urial, Asiatic mouflon and European mouflon) ^1^, 5 □ 6 kya for European mouflon and domestic sheep ^24^, ∼11 kya for Asiatic mouflon and domestic sheep ^4^, and ∼1.26 mya ^1^ for urial and domestic sheep.

### Functional annotation

The genes which overlapping with the concatenated introgressed regions detected by the modified *f*_d_ value were annotated. We annotated and categorized the functions of genes using DAVID v6.8 ^119^ (https://david.ncifcrf.gov/). FDR, Bonferroni and Benjamini-Hochberg adjusted *p*-values were estimated with *p*-value < 0.05 as statistically significant. GO and KEGG pathway enrichment analyses were implemented using DAVIDv6.8 ^119^ (https://david.ncifcrf.gov/).

#### Ethics statement

All animal work was conducted according to a permit (No. IOZ13015) approved by the Committee for Animal Experiments of the Institute of Zoology, Chinese Academy of Sciences (CAS), China. For domestic sheep, animal sampling was also approved by local authorities where the samples were taken.

#### Life Sciences Reporting Summary

Further information on research design is available in the Nature Research Reporting Summary linked to this article.

#### Data availability

Raw sequencing data that support the findings of this study will deposit in the European Nucleotide Archive (ENA) with the corresponding accession codes xxxx and xxxx after acceptance. Source data for Supplementary Figs. 2, 3, 15 are presented in the Supplementary Tables. Additional data such as raw image files and in-house scripts that support this study are available from the first authors upon request.

## ACKNOWLEDGEMENTS

This study was financially supported by grants from the National Key Research and Development Program-Key Projects of International Innovation Cooperation between Governments (2017YFE0117900), the External Cooperation Program of Chinese Academy of Sciences (152111KYSB20190027), the National Natural Science Foundation of China (Nos. 31661143014, 31825024 and 31972527), the Second Tibetan Plateau Scientific Expedition and Research Program (STEP) (No. 2019QZKK0501), and the Taishan Scholars Program of Shandong Province (No. ts201511085). We thank Ming-Shan Wang, Sheng Wang, Hua-Jing Teng, Da-Qi Yu, Peter Wilton, Débora YC Brandt for their technical help with the statistical analysis. We express our thanks to the owners of the sheep for donating samples (see Supplementary Table 1). Thanks are also due to a number of persons for their help during sample collection.

## AUTHOR CONTRIBUTIONS

M.-H.L. conceived the study. M.-H. L. and R.N. supervised the study. Z.-H.C. and Y.-X.X. conducted the laboratory work. Z.-H.C., X.-L.X., G.-J.L. contributed the data analysis. D.-F.W., D.A.G. provided the help for coding. X.-L.X., G.-J.L. performed the analysis of SVs. Z.-H.C., X.-L.X., Y.-X.X., D.W.C., A.E., J.A.L., R.N. and M.-H.L. wrote or revised the paper. K.P., I.A., D.W.C., J. K., M.N., V.R. contributed samples or provided help during the sample collection. All the authors reviewed and approved the final manuscript.

